# HA stability regulates H1N1 influenza virus replication and pathogenicity in mice by modulating type I interferon responses in dendritic cells

**DOI:** 10.1101/744052

**Authors:** Marion Russier, Guohua Yang, Benoit Briard, Victoria Meliopoulos, Sean Cherry, Thirumala-Devi Kanneganti, Stacey Schultz-Cherry, Peter Vogel, Charles J. Russell

## Abstract

Hemagglutinin (HA) stability, or the pH at which the HA is activated to cause membrane fusion, has been associated with the replicative fitness, pathogenicity, transmissibility, and interspecies adaptation of influenza A viruses. Here, we investigated several mechanisms by which a destabilizing HA mutation, Y17H (activation pH 6.0), may attenuate virus replication and pathogenicity in DBA/2 mice, compared to wild-type (WT; activation pH 5.5). Extracellular lung pH was measured to be near neutral (pH 6.9–7.5). WT and Y17H viruses had similar environmental stability at pH 7.0; thus, extracellular inactivation was unlikely to attenuate Y17H virus. The Y17H virus had accelerated single-step replication kinetics in MDCK, A549, and Raw264.7 cells. The destabilizing mutation also increased early infectivity and type I interferon (IFN) responses in mouse bone marrow–derived dendritic cells (DCs). In contrast, the HA-Y17H mutation reduced multistep replication in murine airway mNEC and mTEC cultures and attenuated virus replication, virus spread, severity of infection, and cellular infiltration in the lungs of mice. Normalizing virus infection and weight loss in mice by inoculating them with Y17H virus at a dose 500-fold higher than that of WT virus revealed that the destabilized mutant virus triggered the upregulation of more host genes and increased type I IFN responses and cytokine expression in DBA/2 mouse lungs. Overall, HA destabilization decreased virulence in mice by boosting early infection in DCs, resulting in greater activation of antiviral responses, including type I IFN. These studies reveal HA stability may regulate pathogenicity by modulating IFN responses.

**Importance:** Diverse influenza A viruses circulate in wild aquatic birds, occasionally infecting farm animals. Rarely, an avian- or swine-origin influenza virus adapts to humans and starts a pandemic. Seasonal and many universal influenza vaccines target the HA surface protein, which is a key component of pandemic influenza. Understanding HA properties needed for replication and pathogenicity in mammals may guide response efforts to control influenza. Some antiviral drugs and broadly reactive influenza vaccines that target the HA protein have suffered resistance due to destabilizing HA mutations that do not compromise replicative fitness in cell culture. Here, we show that despite not compromising fitness in standard cell cultures, a destabilizing H1N1 HA stalk mutation greatly diminishes viral replication and pathogenicity in vivo by modulating type I IFN responses. This encourages targeting the HA stalk with antiviral drugs and vaccines as well as reevaluating previous candidates that were susceptible to destabilizing resistance mutations.

## Introduction

Influenza A virus (IAV) isolates vary in fitness and pathogenicity because of differences in the viral genetics and in the ability of the viruses to stimulate or evade host responses. In the H1– H16 subtypes, the influenza hemagglutinin (HA) surface glycoprotein binds sialic acid– terminating moieties on the host-cell plasma membrane (1, 2), is triggered by endosomal low pH to undergo irreversible conformational changes that cause membrane fusion (3), and contributes to virus assembly and morphology (4, 5). Genetic differences, between strains or due to mutations, that alter fundamental properties of the HA protein can alter virus replication, tropism, host range, and pathogenicity (reviewed in references (6, 7)). For example, isolates containing an HA protein with a single Arg residue in its cleavage site are restricted to replication in the lungs where extracellular proteases are available, while isolates encoding the furin protease consensus sequence Arg-X-Lys/Arg-Arg^v^ undergo cleavage maturation intracellularly in the secretory pathway, expanding tropism and increasing pathogenicity in chickens and mammals (8, 9). Mutations in the HA receptor-binding pocket that switch preferential binding from α2,3- to α2,6-linked sialic acid are necessary for avian influenza viruses to adapt to humans (10–12). Two recent studies showed that a mutation that increases HA stability (by decreasing the pH threshold at which the protein is activated) was necessary for gain-of-function airborne transmissibility of H5-subtype viruses in ferrets (13, 14). A stable HA protein was also necessary for airborne transmission of influenza A/H1N1/2009 virus in ferrets and for pandemic potential in humans (15). Numerous mutations that alter HA stability are located in the conserved stalk region (16), which is a target for antiviral drugs and broadly immunogenic vaccines (17). Susceptibility to stalk-based antivirals and vaccines can be altered by changes in HA stability (18, 19). In view of these facts, an understanding of how HA stability regulates IAV replication and pathogenesis bears upon the emergence of pandemic influenza and the development of HA-based control efforts.

During IAV entry into cells, receptor binding at the plasma membrane induces virion internalization (20) via clathrin-dependent or -independent endocytosis (21, 22). Within 5 min, internalized virions are exposed to a pH of 6.0–6.5 in early endosomes, and over a period of 30– 40 min or more they are exposed to pHs of 5.0–5.5 in late endosomes and 4.6–5.0 in lysosomes (23, 24). At a threshold pH, HA trimers are triggered to undergo irreversible conformational changes that cause membrane fusion (reviewed in reference (3)). The threshold pH for HA activation (and subsequent inactivation) ranges from approximately 4.8 to 6.2 (16, 25, 26). In general, avian IAVs have less stable HA proteins with higher activation pH values and human-adapted IAVs have more stable HA proteins. Less stable avian-origin HA proteins can enable virus replication in macrophages (27) and allow virus entry into early endosomes, thereby avoiding virion interactions with the IFITM2 and IFITM3 proteins that stimulate interferon (IFN)-induced antiviral activity (28).

Dozens of mutations throughout the HA trimer have been identified as altering the HA-activation pH, and many of these mutations are associated with adaptations (reviewed in references (16, 29, 30)). Destabilizing mutations enhance virus replication in the presence of ammonium chloride and high concentrations of amantadine, which raise the endosomal pH (31–34). The replication of A/PR/8/34 (H1N1) virus, which has an HA activation pH of 5.0–5.1 (26), is enhanced in Vero cells by a destabilizing mutation that increases the activation pH by 0.2 pH units (35). NS1-deleted vaccine candidates with HA-activation pH values of 5.5–5.8 have infectivity and immunogenicity higher than those of related variants triggered at pH 6.0–6.3 (36, 37). In H5 viruses with relatively unstable HA proteins, stabilizing mutations have been shown to increase upper respiratory tract replication in mice and ferrets (38–40) and to enable gain-of-function transmissibility in ferrets (13, 14); however, these mutations decrease the replication, virulence, and transmissibility in avian species (41–43).

Adaptations of IAVs to mice suggest that there is an optimal HA-activation pH range of approximately 5.4–5.8 for replication in the lungs. In the context of 20^th^-century seasonal H1N1 and H3N2 viruses that are activated at pH 5.2–5.3, adaptation to mouse lungs yielded destabilization mutants that elevated the HA-activation pH to 5.6–5.8 (44–46). However, the adapted viruses also exhibited changes in receptor binding. For H5N1 and A/2009/H1N1 viruses containing point mutations that did not alter receptor-binding specificity or avidity, HA proteins that were activated at pH 5.4–5.5 boosted replication in mouse lungs and increased pathogenicity when compared to those activated at pH 5.9–6.3 (15, 39).

The mechanisms by which HA stability regulates influenza virus replication, pathogenesis, transmissibility, and host range are unclear. As viruses containing unstable HA proteins (with activation pH > 5.8) have less environmental stability (41), we hypothesized that a relatively stable HA protein was required for maximum in vivo infectivity and replication to avoid extracellular inactivation in the respiratory tract (16). To test this hypothesis, we studied infection with A/TN/1-560/2009 (H1N1), a 2009 pandemic virus, and two viruses with HA stability–altering mutations (Y17H and R106K). The wild-type (WT) HA protein is activated at pH 5.5, whereas a Y17H mutation in the HA1 fusion-peptide pocket increases the activation pH to 6.0 and an R106K mutation in the HA2 coiled-coil core decreases the activation pH to 5.3 (15). Neither mutation altered HA protein expression, cleavage, maturation, receptor-binding avidity, or receptor-binding specificity. Both mutant viruses exhibited multistep replication kinetics similar to those of WT virus in MDCK cells; however, Y17H virus had reduced replication and was less lethal than WT virus in mice (15). The objective of this study was to use a mouse model to determine the mechanism by which HA stability regulates A/H1N1/2009 replication and pathogenicity.

## Materials and Methods

### Cells

The following cell lines were obtained from the American Type Culture Collection (ATCC): Madin-Darby canine kidney (MDCK) (ATCC CCL-34); murine lung adenoma LA-4 (ATCC CCL-196); human lung epithelial carcinoma A549 (ATCC CCL-185); and murine macrophage, Abelson murine leukemia virus–transformed, Raw264.7 (ATCC TIB-71). MDCK cells were propagated in culture in Dulbecco’s modified Eagle’s medium (MEM) supplemented with 5% fetal bovine serum (FBS) and 1% penicillin-streptomycin (Pen-Strep) at 37 °C in 5% CO_2_. Both LA-4 and A549 cells were propagated in culture in Kaign’s modification of Ham’s F-12 medium with L-glutamine (F-12K) supplemented with 10% FBS and 1% Pen-Strep at 37 °C in 5% CO_2_. Raw264.7 cells were propagated in culture in RPMI-1640 medium supplemented with 10% FBS and 1% Pen-Strep at 37 °C in 5% CO_2_. To prepare and propagate murine nasal epithelial cells (mNECs) and murine tracheal epithelial cells (mTECs), nasal turbinate and trachea were collected and cells isolated using collagenase (200 U/ml)/dispase (2.5 U/ml) in F12 containing Pen-Strep or 0.15% filter-sterilized pronase diluted in F-12 containing Pen-Strep respectively. After appropriate incubation, cells were pelleted by centrifugation, washed, and resuspended in ammonium chloride potassium (ACK) lysis buffer to remove red blood cells. Following resuspension in media, cells were pelleted by centrifugation and plated at 3.3 x 10^4^ cells per 0.32 cm^2^ transwell inserts pre-coated with rat tail collagen and grown in DMEM-F12 supplemented with 200 mM glutaMAX, 7.5% sodium bicarbonate, insulin (10 ug/ml), transferrin (5 ug/ml), cholera toxin (100 ng/ml), murine epidermal growth factor (25 ng/ml), bovine pituitary extract (30 ug/ml), 5% fetal bovine serum, retinoic acid (5 x 10^-10^ M), and Pen-Strep until confluency. Once confluent, apical media was removed and basal media was replaced with ALI media (DMEM-F12 containing 200 mM glutaMAX, 7.5% sodium bicarbonate, Pen-Strep, 2% NuSerum and retinoic acid). Cells were used after full differentiation (TER > 1000), which was typically after one week for mNEC and 1 to 3 weeks for mTEC. Bone marrow-derived macrophages (BMDMs) and dendritic cells (BMDCs) were prepared as described previously (47). Briefly, primary BMDMs were grown for 6 days in IMDM (12440-053, Gibco) supplemented with 1% non-essential amino acids (Gibco), 10% FBS (S1620, lot number 221C16, BioWest), 30% medium conditioned by L929 mouse fibroblasts, and 1% penicillin–streptomycin (Gibco). Primary BMDCs were grown in RPMI (10–040-CV, cellgro, Corning) supplemented with 10% FBS (S1620, lot number 221C16, BioWest), 1% non-essential amino acids (Gibco), 1% sodium pyruvate (Gibco), 1% penicillin–streptomycin (Gibco), 0.1% β-mercaptoethanol, and 20 ng.ml^−1^ granulocyte–macrophage colony-stimulating factor for 7 days. BMDMs and BMDCs were seeded in antibiotic-free media at a concentration of 1 x 10^6^ and 3 x 10^6^ cells respectively onto 12-well plates.

### Viruses

Influenza A/Tennessee/1-560/2009 (H1N1) WT, Y17H, and R106K viruses were previously generated by reverse genetics, sequenced, and characterized (15, 48). The Y17H mutation is located in HA1 at residue 17 in H3 numbering, position 7 from the N-terminal cleavage site (49), and at residue 24 in the full-length H1 sequence. The R106K mutation is at HA2 position 106 in both H1 and H3 numbering.

### Mice

Six-week-old female DBA/2J mice were obtained from The Jackson Laboratory. MAVS^-/-^, and Myd88^-/-^ TRIF^-/-^ mice were described previously (50, 51). All mice were backcrossed to the C57BL/6 background and bred in house.

### Animal ethics statement

Animal experiments were conducted in an ABSL2+ facility, using procedures compliant with NIH requirements and with the Animal Welfare Act. The St. Jude Animal Care and Use Committee reviewed and approved the animal experiments (protocol numbers 459 and 464).

### Animal experiments

Seven-week-old DBA/2J mice were anesthetized with isoflurane and inoculated intranasally with virus in 30 µL of PBS. Clinical symptoms (including weight loss and mortality) were monitored daily. To enable the collection of respiratory tissues, groups of mice were euthanized with CO_2_ after 2 and 5 days of infection. Bronchoalveolar lavage fluid (BALF) was collected by washing the lungs three times with 0.5 mL of PBS containing 2 mM EDTA (for a total volume of 1.5 mL) via a catheter. The BALF was centrifuged, then the total number of infiltrating cells was counted by flow cytometry and the supernatants were stored at −80 °C. Lung tissues were collected for use in the various assays described below.

### Virus titrations and environmental persistence assays

At 2 and 5 days post infection (DPI), mice were euthanized and nasal turbinates, tracheae, and lungs were collected. After the tissues had been homogenized using a Qiagen Tissue Lyser II (30/s frequency for 30s, operated twice), the homogenates were clarified by centrifugation at 8,000 RPM for 15 min at 4 °C in an Eppendorf 5417R centrifuge. The infectivity of the supernatants was titrated by 50% tissue culture infective dose (TCID_50_) assays. One day before the virus infection, 3–5 × 10^4^ MDCK cells in culture medium were dispensed into each well of a 96-well tissue-culture plate and incubated at 37 °C in 5% CO_2_. On the day of infection, tissue supernatants were serially diluted 10-fold in virus infection medium supplemented with 1 µg/mL L-1-tosylamide-2-phenylethyl chloromethyl ketone (TPCK)-trypsin (Worthington Biochemical Corporation, Lakewood, NJ). The plates were rinsed twice with PBS, and 200 µL of diluted tissue supernatant sample was added to each well containing MDCK cells. The plates were incubated at 37 °C in 5% CO_2_ for 3 days, then 50 µL of the culture from each well was transferred to a new 96-well round-bottom plate, then 50 µL of 0.5% turkey red blood cells in PBS was added to each well. The plates were incubated for 30 min at room temperature. The highest dilution that tested positive for virus was recorded. TCID_50_ values were calculated using the method of Reed and Muench (52).

To measure virus persistence, thawed virus stocks were diluted to approximately 10^6^ TCID_50_ /mL in pH-adjusted PBS (pH 6.4 or 7.0). Viruses were incubated at 37 °C and collected at the specified time points to measure the residual infectivity by TCID_50_ assay. The time required for a 10-fold decrease in virus titer (Rt) was calculated by least-squares linear regression of values above the limit of detection, using GraphPad Prism 7.

### Lung pH measurements

A pH-1 micro fiber-optic pH transmitter and a pH-Microsensor (a needle-type fiber-optic microsensor) were purchased from PreSens Precision Sensing GmbH (Regensburg, Germany). The microsensor was calibrated manually according to the manufacturer’s manual before experimental measurements. In brief, calibration details listed on the Final Inspection Protocol of the microsensors, which were provided together with each sensor, were typed into the pH1-View, Manual Calibration under Calibration. DBA/2J mice were inoculated intranasally with 30 μL of PBS or virus inoculum (750 or 375,000 PFU). At 2 and 5 DPI, mice were euthanized, their chests were opened immediately, and the pH Microsensor was inserted into their lungs. In each case, the pH was recorded 30 s after probe insertion.

### Virus replication curves

For multistep virus growth-curve experiments, MDCK, LA-4, mNEC, and mTEC cells were infected with virus at a multiplicity of infection (MOI) of 0.01 PFU/cell and incubated at the indicated temperatures (33 °C, 37 °C, or 39.5 °C) in 5% CO_2_ for 1 h, after which they were rinsed three times with PBS. Infection medium was then added, and the cells were incubated at the indicated temperatures (33 °C, 37 °C, or 39.5 °C) in 5% CO_2_. Supernatants were collected at the indicated time points, stored at −80 °C, and titrated in MDCK cells by TCID_50_ assay. For single-step growth-curve experiments, MDCK, A549, and Raw264.7 cells were infected with virus at an MOI of 3 PFU/cell for 1 h at 37 °C, after which the infected cells were rinsed once with saline buffer (pH 2.2) and twice with PBS. The cells were incubated at 37 °C in 5% CO_2_ until harvested at the specified time points. Samples were stored at −80 °C until they could be titrated in MDCK cells by TCID_50_ assay.

### Histopathology

The lungs from control and virus-infected mice were fixed via intratracheal infusion with and then immersion in 10% buffered formalin solution. Tissues were paraffin embedded, sectioned, and immunohistochemical labeling of viral antigen was completed by using a primary goat polyclonal antibody (US Biological, Swampscott, MA) against influenza A, USSR (H1N1) at 1:1000 and a secondary biotinylated donkey anti-goat antibody (catalog number sc-2042; Santa Cruz Biotechnology, Santa Cruz, CA) at 1:200 on tissue sections subjected to antigen retrieval for 30 minutes at 98°C. The extent of virus spread was quantified by first capturing digital images of whole-lung sections stained for viral antigen by using an Aperio ScanScope XT Slide Scanner (Aperio Technologies, Vista, CA) then manually outlining fields with the alveolar areas containing virus antigen-positive pneumocytes highlighted in red (defined as ACTIVE infection). Those lesioned areas containing minimal or no antigen positive cells were highlighted in yellow (defined as INACTIVE infection). The percentage of each lung field with infection/lesions was calculated using the Aperio ImageScope software. In addition, a pathologist “blinded” to treatment group identity evaluated pulmonary lesions in HE-stained histologic sections and assigned scores based on their severity and extent as follows: 0 = no lesions; 1 = minimal, focal to multifocal, barely detectable; 15 = mild, multifocal, small but conspicuous; 40 = moderate, multifocal, prominent; 80 = marked, multifocal coalescing, lobar; 100 = severe, diffuse, with extensive disruption of normal architecture and function.

### Chemokine and cytokine assays

IFNβ was measured with an enzyme-linked immunosorbent assay (ELISA) kit (BioLegend). Other chemokines and cytokines were measured by the MILLIPLEX mouse magnetic bead assay (Millipore) according to the manufacturer’s instructions.

### Microarray analyses

Pieces of lung were preserved in RNAlater (Ambion), and lung tissues were homogenized using a Qiagen Tissue Lyser II. RNA was extracted from lung samples with an RNeasy Kit (Qiagen), and microarray analyses were performed by the Hartwell Center for Bioinformatics and Biotechnology at St. Jude Children’s Research Hospital. Briefly, total RNA (100 ng) was converted to biotin-labeled cRNA with an Ambion Wild-Type Expression Kit (Life Technologies) and was hybridized to a Mouse Gene 2.0 ST GeneChip microarray (Affymetrix). After the microarray was stained and washed, the array signals were normalized and transformed into log2 transcript expression values with the robust multi-array average algorithm (Partek Genomics Suite version 6.6). Differential expression was defined by applying a difference in expression of 0.5-fold (log2 signal) between conditions. Lists of differentially expressed transcripts were analyzed for functional enrichment with the Gene Ontology Resource at geneontology.org (53, 54). Visualization of gene expression profile (heat map creation) was done using TIBCO Spotfire software, version 10.1.0.

### Real-time quantitative RT-PCR

RNAlater-preserved tissues were homogenized, and total RNA was extracted using the RNeasy Mini Kit (Qiagen). The levels of virus-specific vRNA, cRNA, and mRNA, as well as GAPDH mRNA, were analyzed by semiquantitative real-time PCR analysis on an Applied Biosystems 7500 Fast Real-Time PCR System (Applied Biosystems, Waltham, MA) as previously described (55). Briefly, total RNA was reverse transcribed using virus-specific RT primers or oligo-dT primers and the SuperScript III First-Strand Synthesis System (Invitrogen, Carlsbad, CA). Real-time quantitative PCR was performed on the Applied Biosystems 7500 system with QuantiTect SYBR Green PCR Master Mix (Qiagen) and the appropriate primers. After normalization of the results to the GAPDH expression levels, the fold-change ratio of expression in virus-infected samples to that in control samples was calculated for each gene by using the ΔΔCt method and was expressed as 2−ΔΔCt.

### Immunoblot analysis

Lung tissues or bone marrow–derived DCs were lysed in RIPA lysis buffer and sample loading buffer. Proteins were separated by electrophoresis on 8%–12% polyacrylamide gels. After the proteins were electrotransferred onto PVDF membranes, nonspecific binding was blocked by incubating the membranes with 5% skim milk or BSA. The membranes were then incubated with primary antibody (anti-pSTAT1 Y701 [clone D4A7; Cell Signaling Technologies; 1:100 dilution], anti-STAT1 p84/p91 [clone E23; Santa Cruz Biotechnology; 1:1,000 dilution], anti-M1 [GeneTex, GTX127356; 1:1,000 dilution], or anti-actin [Santa Cruz Biotechnology, SC-47778; 1:10,000 dilution]) and then with secondary antibody (horseradish peroxidase–conjugated anti-rabbit IgG [Cell Signaling Technologies, 7074S] or anti-mouse IgG [Cell Signaling Technologies, 7076S], both at a 1:5,000 dilution]).

### Immunofluorescence microscopy

Bone marrow–derived DCs were infected as previously described and seeded in a 96-well plate. At 4 h post infection, cells were fixed in PBS–4% paraformaldehyde for 15 min. They were then permeabilized with PBS–0.5% Triton X-100 for 10 min, blocked in PBS–10% FBS for 1 h, and stained with an antibody to influenza A NP (Millipore, MAB8257) overnight at 4 °C. After the plate had been washed, it was incubated with anti-mouse IgG–Alexa 488 secondary antibody (Invitrogen) in 10% FBS in PBS for 1 h. The plate was then imaged in a fluorescence multiplate reader (Synergy 2 Multi-Mode Microplate Reader; BioTek).

### Statistical analyses

GraphPad Prism 7.03 software was used for data analysis. Statistical significance for two groups was determined by Student’s *t*-test (two-tailed). Statistical significance for three or more groups was determined by one-way ANOVA followed by Tukey’s multiple comparisons test. We considered *P* values of less than 0.05 to indicate statistical significance.

## Results

### Y17H virus is attenuated for infectivity, replication, and virulence in mice

The HA protein of A/Tennessee/1-560/2009 (H1N1) WT virus was previously shown to be activated for membrane fusion or, in the absence of target cells, inactivated by low-pH buffer at a midpoint pH of 5.5. HA2 stalk mutations Y17H and R106K altered HA stability to pH 6.0 and 5.3, respectively, yet these mutated proteins retained similar expression levels, cleavage, and receptor-binding specificity (15, 48). To investigate the mechanisms by which HA stability alters infectivity and pathogenicity, we inoculated groups of DBA/2 mice intranasally with various doses of reverse-genetics (r.g.) WT, Y17H, and R106K viruses. The mouse infectious dose 50 (MID_50_) of these viruses decreased with decreasing HA activation pH (Table 1); thus, increased HA stability was associated with increased infectivity. WT virus had an MLD_50_ value of 11,000 PFU; the R106K mutation increased the MLD_50_ to 20,100 PFU; and 80% of Y17H virus– infected mice survived infection with 375,000 PFU, the highest dose tested (Table 1). At equivalent doses, WT and R106K viruses induced similar weight loss, whereas Y17H virus caused substantially less weight loss (Figure 1A–C). For example, at a dose of 750 PFU, mice in the WT- and R106K virus–infected groups exhibited approximately 10% weight loss, whereas mice in the Y17H virus–infected group lost only approximately 2% of their weight. We euthanized additional groups of mice at 2 and 5 DPI and measured the viral loads. WT and R106K viruses yielded similar titers (*P* > 0.05) at all inoculated doses except for 750 PFU in trachea at 2 DPI (Figure 1D-–I). As infection with R106K virus yielded viral loads and weight loss similar to those observed with WT virus, R106K virus was excluded from subsequent mechanistic studies. Titers of Y17H virus were only 10% of WT virus for many of the inoculation doses used (Figure 1D–I). The maximal nasal titer of WT virus at 2 DPI was approximately 10^5^ TCID_50_/mL, whereas that of Y17H virus was lower by a factor of approximately 100 (Figure 1D). At 2 DPI with a 750-PFU dose, WT virus reached its maximal lung titer of approximately 10^6^ TCID_50_/mL, whereas the corresponding lung titer of Y17H virus was lower by a factor of approximately 200. Overall, these studies showed that the HA-stabilizing mutation R106K had little effect on pathogenicity, whereas the HA-destabilizing Y17H mutation was highly attenuating.

**Figure 1.**
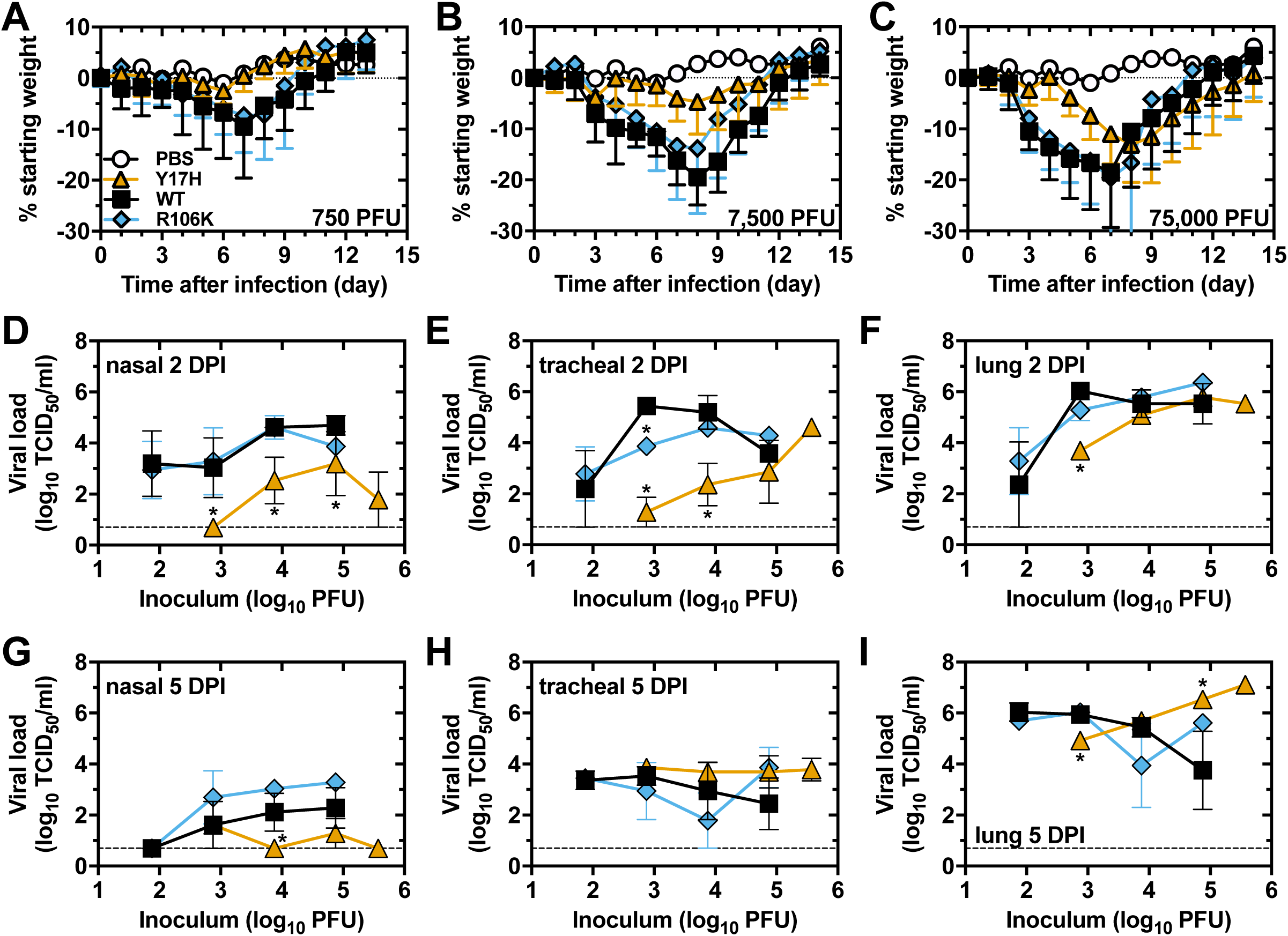
Weight loss and viral tissue titers in mice. **(A–C)** Changes in starting weight of DBA/2 mice after inoculation with 750 (A), 7500 (B), or 75,000 (C) PFU of virus. Values are the combined mean (± SD) from two independent experiments with 10 mice total. **(D–I)** Nasal, tracheal, and lung tissue titers at 2 DPI (D–F) or 5 DPI (G–I). Inoculated doses were 75, 750, 7500, 75,000, or 375,000 PFU. Y17H virus was not inoculated at a dose of 75 PFU, and WT and R106K viruses were not inoculated at a dose of 375,000 PFU. Values are the mean (± SD) from three mice. The legend in panel A shows the symbols used in all panels. Black squares: WT virus; blue diamonds: R106 virus; orange triangles: Y17H virus; white circles: PBS mock infection. For panels D–I, the statistical significance was determined by Student’s *t*-test, comparing each set of results to those for WT virus at the equivalent dose: * *P* < 0.05.

**Table 1.** Infectious and lethal dose values in mice compared to HA activation pH values

| Virus | Activation pH<sup>a</sup> | MID50 (PFU)<sup>b</sup> | MLD<sub>50</sub> (PFU)<sup>c</sup> |
| --- | --- | --- | --- |
| Y17H | 6.0 | 48 | >375,000 |
| WT | 5.5 | 4.7 | 11,000 |
| R106K | 5.3 | 0.68 | 20,100 |

<sup>a</sup>HA activation pH values measured by syncytia formation assay and pH of inactivation of virus infectivity assay, both of which were in numerical agreement.
<sup>b</sup>Mouse infectious dose 50 value calculated by the method of Reed and Muench.
<sup>c</sup>Mouse lethal dose 50 value calculated by the method of Reed and Muench. For the Y17H group, 80% of mice survived infection after inoculation with a dose of 375,000 PFU, the highest dose tested.

### Reference and normalized inoculation doses of WT and Y17H viruses

For subsequent in vivo studies, we selected 750 PFU as the reference dose. This dose yielded a 10-fold difference in weight loss and 10-to 100-fold differences in lung viral titers (Figure 1), and it had previously resulted in differences in histopathology, cellular infiltration, and cytokine/chemokine expression (15). To normalize the viral titers and weight loss for subsequent studies of host responses, we considered two additional groups. A 75-PFU inoculum of WT virus resulted in an average lung viral titer that was lower by a factor of approximately 10 at 2 DPI (*P* = 0.4731, non-significant) but 10-fold higher at 5 DPI (*P* = 0.0814, non-significant) (Figure S1A) than the corresponding titer obtained with a 10-fold higher 750-PFU inoculum of Y17H virus at those time points. With respect to weight loss, a 75-PFU dose of WT virus yielded a maximum loss of approximately 9%, whereas a 750-PFU dose of Y17H virus caused significantly less weight loss (*P* < 0.05 on days 7–10) (Figure S1C). Thus, a 75-PFU dose of WT virus did not result in pathogenicity like that caused by a 10-fold higher 750-PFU dose of Y17H virus. We next inoculated mice with a 500-fold higher dose (375,000 PFU) of Y17H virus and compared the results to those in mice inoculated with 750 PFU of WT virus. These two groups of mice exhibited similar weight loss (*P* > 0.05) (Figure S1D). Compared to a 750-PFU dose of WT virus, a 375,000-PFU dose of Y17H virus resulted in lung titers approximately 0.5-log lower at 2 DPI (*P* = 0.2193) and approximately 1-log higher at 5 DPI (*P* = 0.0114) (Figure S1B). Overall, the data showed that increasing the dose of Y17H virus to 375,000 PFU resulted in similar weight loss and comparable, albeit unequal, lung viral titers to those seen with a dose of 750 PFU of WT virus. Thus, in addition to the groups of mice infected with 750 PFU of WT or Y17H viruses, we included a group infected with 375,000 PFU of Y17H virus in our subsequent analyses of potential differences in the host responses.

### Y17H virus retains WT-like infectivity at the extracellular pH of murine lungs

One mechanism by which the HA-destabilizing Y17H mutation could attenuate virus growth in vivo is by extracellular inactivation if the respiratory tract is mildly acidic (16). To measure the extracellular pH in murine lungs, groups of mice were inoculated intranasally with WT virus (750 PFU), Y17H virus (750 or 375,000 PFU), or PBS. In PBS-control mice, the average lung pH was 7.04 (Figure 2A). The average lung pH in infected mice at 2 and 5 DPI was increased to approximately 7.3–7.4 (*P* < 0.02 or less compared to PBS) for all groups except mice infected for 5 days with 750 PFU of Y17H virus, in which the lung pH recovered to 7.04. By 35 DPI, the lung pH in all groups had recovered to approximately 7.0–7.1. Overall, this experiment showed that the lungs of DBA/2 mice remain at a near-neutral pH whether they are uninfected, are infected with a pH1N1 virus, or have recovered from an infection.

**Figure 2.**
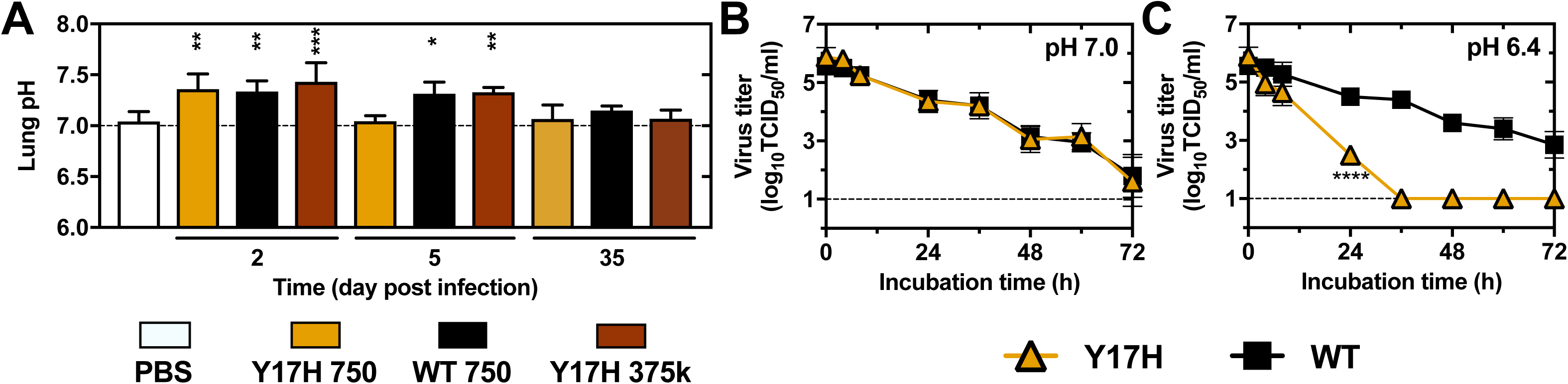
Murine extracellular lung pH and virion environmental stability. **(A)** Lung pH values in DBA/2 mice. Mice were inoculated intranasally with PBS (white), 750 PFU of WT virus (black), 750 PFU of Y17H virus (orange), or 375,000 PFU of Y17H virus (brown). After 2, 5, or 35 days of infection, extracellular pH values were measured in the lungs. Reported values are means (n = 6) with standard deviations. **(B,C)** Environmental stabilities of WT and Y17H viruses. Virus stocks were thawed and diluted to approximately 10^6^ TCID_50_/mL in PBS with the pH adjusted to 7.0 (B) or 6.4 (C) then incubated for the specified times at 37 °C before the residual infectivity was measured by TCID_50_ assay. Groups included those infected with WT virus (black squares) or Y17H virus (orange triangles). For panels B and C, the values are reported as means (n = 3) with standard deviations. Statistical significance tests compared the results to those for PBS (panel A) or WT virus (panels B,C): * *P* < 0.05, ** *P* < 0.01, *** *P* < 0.001, **** *P* < 0.0001.

To measure the persistence of virion infectivity (environmental stability) at neutral pH, we incubated aliquots of WT and Y17H viruses at 37 °C in buffer of pH 7.0 or 6.4 and measured the residual infectivity as a function of time by TCID_50_ assay (Figure 2B,C). At pH 7.0, the titers of both viruses dropped from approximately 5.7 log_10_ to 1.7 log_10_ TCID_50_/mL over a period of 72 h. When virus aliquots were incubated at pH 6.4, the titer of Y17H virus decayed at an accelerated rate compared to that of WT virus. The 90% reduction time (Rt value) is the time required for a 90% (1 log_10_) decrease in the viral titer (56). At pH 7.0, the Rt values for the WT and Y17H viruses were similar (21.7 ± 1.5 h and 20.4 ± 2.1 h, respectively); however, at pH 6.4, the Rt values for the WT and Y17H viruses were 26.6 ± 1.8 h and 7.4 ± 0.6 h, respectively. In summary, Y17H virions were inactivated at a rate 3.6 times faster than WT virions when exposed to pH 6.4 but were inactivated at a similar rate to WT virions when exposed to pH 7.0. As the measured pH of murine lungs did not drop below pH 7.0, a value at which the environmental stability of Y17H virus showed little to no accelerated decay, we concluded that the attenuation of Y17H virus in murine lungs was not due to extracellular inactivation.

### Replication in cultured cells

We previously found that Y17H virus exhibited multistep replication kinetics like those of WT virus in MDCK, A549, and normal human differentiated epithelial (NHBE) cells at 37 °C (15). Here, we performed additional single- and multistep growth-curve experiments. In MDCK cells inoculated at an MOI of 0.01 PFU/cell and incubated at 33 or 39.5 °C, Y17H virus grew in a similar manner to WT virus (Figure 3A,B). Y17H and WT viruses also had similar replication kinetics (multistep and single-step) in LA-4 murine lung adenoma epithelial cells (Figure 3C,F). In differentiated murine nasal (mNEC) and tracheal (mTEC) epithelial cells grown at an air-liquid interface, replication of Y17H virus was delayed by approximately 12–24 h (Figure 3E,F). Additionally, the maximal growth of Y17H virus in mTEC cells was less than 10% of that in MDCK cells. In Raw264.7 mouse macrophage cells, WT virus did not produce an infection, whereas Y17H virus replicated (Figure 3I). For completeness, we performed single-step growth-curve experiments using MDCK and A549 cells. Surprisingly, the data showed that the growth of Y17H virus was enhanced relative to that of WT virus (Figure 3G,H). In summary, Y17H virus replicated faster than WT virus in single-step growth-curve experiments in MDCK, A549, and Raw264.7 cells; Y17H virus exhibited similar growth to WT virus in multistep growth-curve experiments in MDCK, A549, and LA-4 cells; but the growth of Y17H virus was delayed in comparison to that of WT virus in mNEC and mTEC cells.

**Figure 3.**
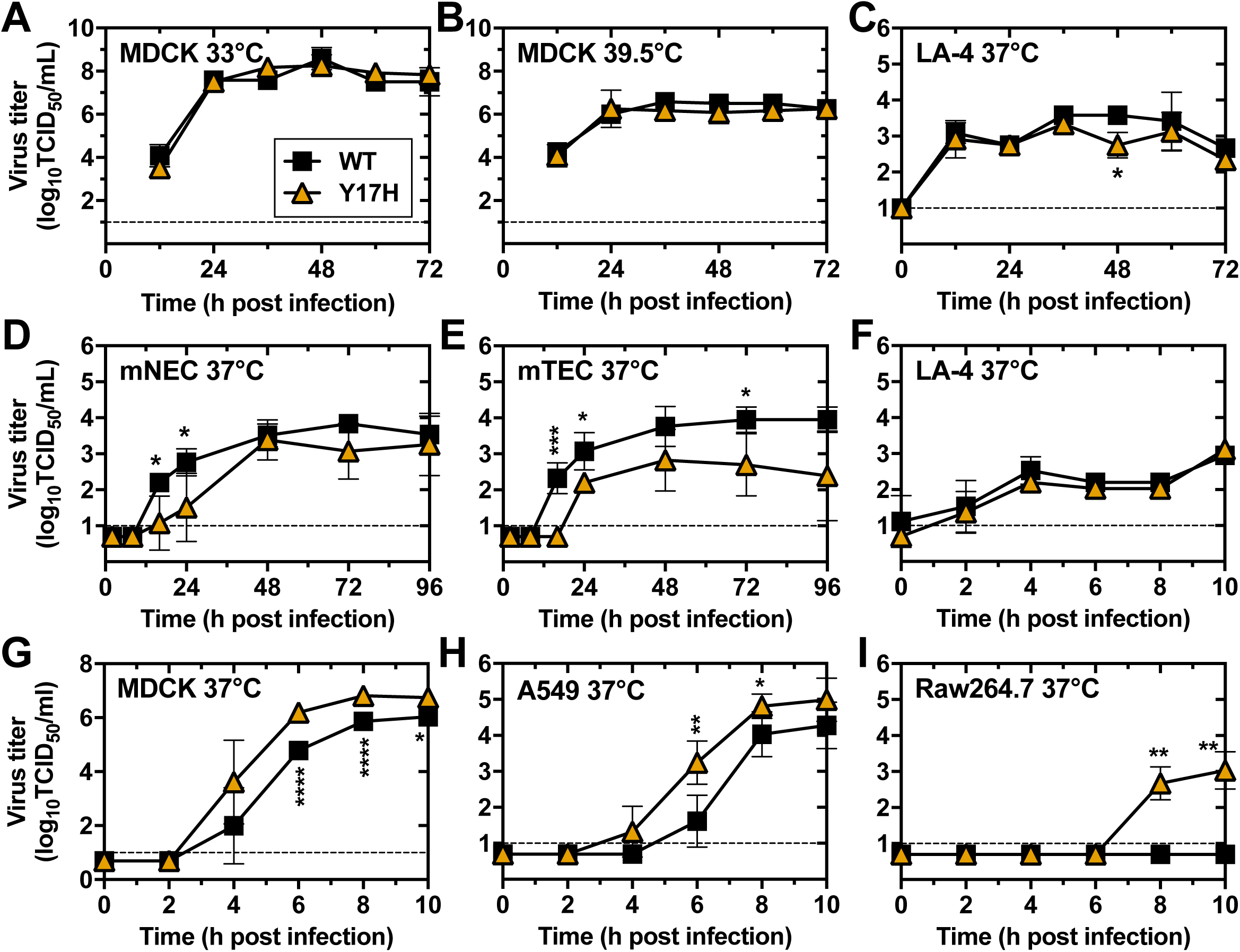
Virus growth in cell cultures. **(A,B)** Multiple-step virus replication (MOI 0.01 PFU/cell) in MDCK cells in culture at 33 °C (A) or 39.5 °C (B). **(C)** Multiple-step virus replication in LA-4 mouse lung adenoma cells in culture at 37 °C. **(D,E)** Multiple-step virus replication in immortalized murine epithelial cells in culture at an air-liquid interface at 37 °C. The differentiated nasal mNEC (D) and tracheal mTEC (E) cells were grown on transwell plates with an air-liquid interface. **(F)** Single-step virus replication in LA-4 mouse lung adenoma cells in culture at 37 °C. **(G)** Single-step virus replication in MDCK cells in culture at 37 °C. **(H)** Single-step virus replication in A549 cells in culture at 37 °C. **(I)** Multiple-step virus replication in Raw264.7 murine macrophage cells in culture at 37 °C. Virus titers are given as the mean (± SD) for three or four replicates of infection with WT virus (black squares) and Y17H virus (orange triangles). For single-step growth curves, cells were infected at an MOI of 3 PFU/cell, and for multiple-step growth curves, cells were infected at an MOI of 0.01 PFU/cell. Statistical significance tests compared the results to those for WT virus: * *P* < 0.05, ** *P* < 0.01, *** *P* < 0.001.

### Pathogenicity at equivalent and normalized doses

To investigate how the HA-destabilizing Y17H mutation affected in vivo spread and pathogenesis, we inoculated groups of five mice intranasally with PBS, WT virus (750 PFU), or Y17H virus (750 or 375,000 PFU) and recovered the lungs at 2 and 5 DPI. Compared to infection with WT virus, infection with Y17H virus at the same dose (750 PFU) resulted in significantly less spread of infection, growth of virus, severity of infection, cellular infiltration, and IL6 and IP10 expression in the lungs (Figures 4 and S2). Thus, Y17H virus was highly attenuated in vivo, recapitulating in vitro replication in mNECs and mTECs, the only differentiated cells used for growth curves. We also wished to compare the results of a 750-PFU infection with WT virus and a 375,000-PFU infection with Y17H virus, because in a previous experiment, these two virus exposures induced similar weight loss and the viruses grew to similar lung titers (Figure S1D). By 5 DPI, infection in the 375,000-PFU Y17H group covered 14% less area, grew to titers approximately 1 log higher, induced approximately 19% more cellular infiltration, and was scored as approximately 17% more severe when compared to infection in the 750-PFU WT group. However, the 750-PFU WT and 375,000-PFU Y17H groups did not differ significantly (*P* > 0.05) at 2 DPI or 5 DPI with respect to virus growth, spread of infection, severity of infection, cellular infiltration, and IL6 and IP10 expression in the lungs (Figures 4 and S2). Thus, inoculating mice with Y17H virus at a dose 500-fold higher than that used for WT virus induced an infection that was sufficiently robust to overcome the attenuation of Y17H virus. This suggested that normalizing Y17H infection with a dose 500-fold higher than the WT virus dose could be used to investigate differential triggering and/or counteracting of antiviral host responses by the two viruses.

**Figure 4.**
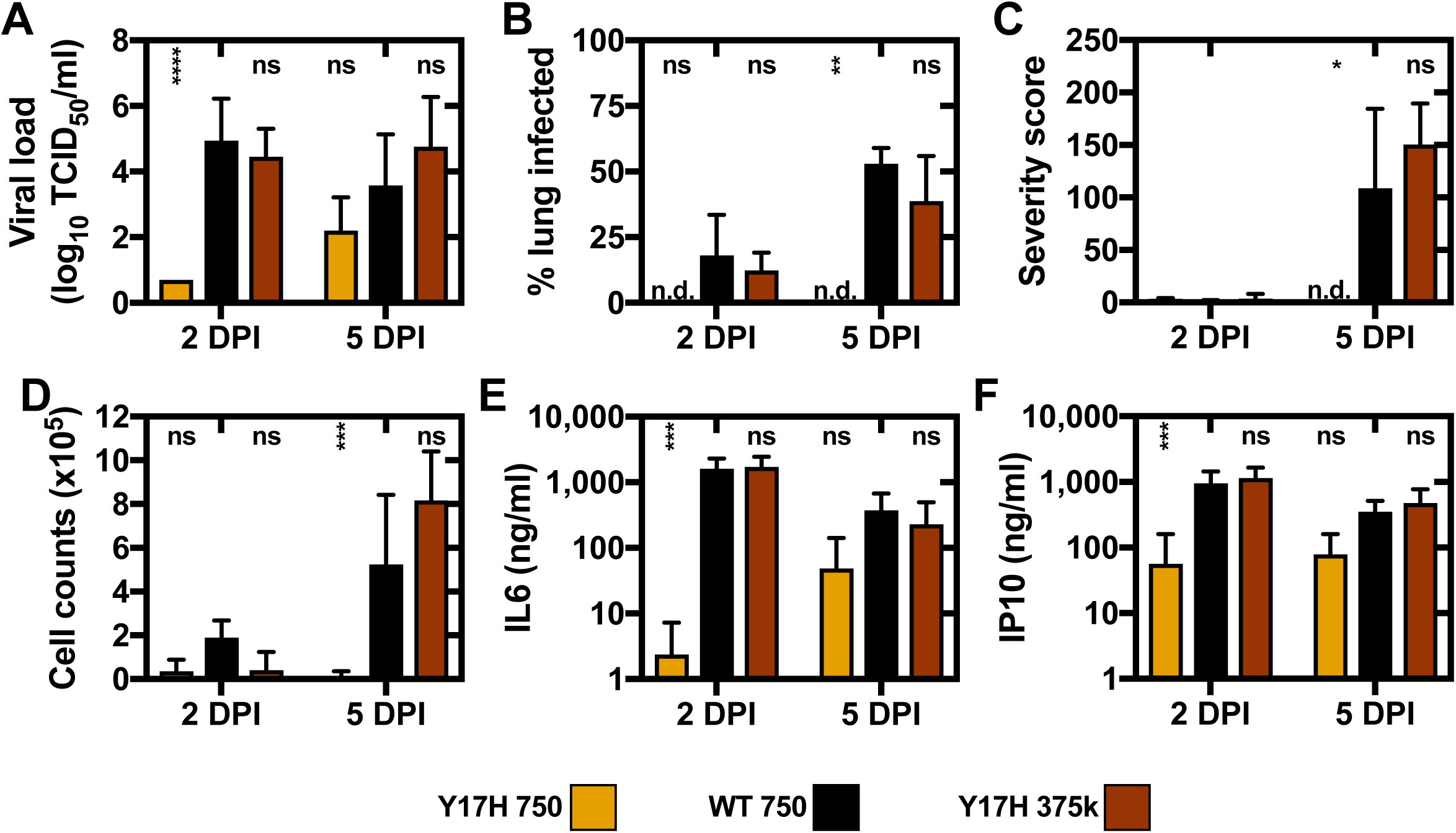
Infection, histopathology, infiltration, and cytokine induction in lungs. Groups of mice were infected with 750 PFU of Y17H virus (orange bars), 750 PFU of WT virus (black bars), or 375,000 PFU of Y17H virus (brown bars). After 2 and 5 days of infection, lungs were recovered to assess the spread of infection, histopathology, cellular infiltration in BALF, and cytokine expression in BALF. **(A)** Lung homogenates were plaque titrated by TCID_50_ assay on MDCK cells. **(B,C)** Lungs were fixed, sectioned, and analyzed microscopically to determine the percentage of cells infected (B) and the severity of the infection (C). **(D)** Cellular infiltration in BALF was measured by flow cytometry. **(E,F)** Expression of IL6 (E) and IP10 (F) in BALF was measured by ELISA. Undetectable values are labeled n.d. Displayed values are the mean (± SD) from five mice per group. Multiple comparisons to the 750-PFU WT group at each time point were analyzed by two-way ANOVA (time and group) followed by Tukey’s post hoc test. Statistical significance: * *P* < 0.05, ** *P* < 0.01, *** *P* < 0.001, **** *P* < 0.0001. ns = not significant (*P* > 0.05).

### Impact of HA stability on antiviral responses

To analyze gene expression to examine if the Y17H virus differentially induces host responses, we inoculated groups of three mice with PBS, WT virus (750 PFU), or Y17H virus (750 or 375,000 PFU). At 2 and 5 DPI, we collected lungs and processed the samples for Affymetrix microarray analysis. At an equivalent dose of 750 PFU, infection with Y17H virus resulted in 21% fewer genes (127 vs. 161) being upregulated than with WT virus after 2 days and 54% fewer genes (142 vs. 308) being upregulated after 5 days (Figure 5A). The upregulation of fewer genes by infection with Y17H virus was consistent with prior experiments showing less virus growth and cytokine expression with the attenuated virus (Figure 4). Despite mice in the 375,000-PFU Y17H group showing virus spread, infiltration, and cytokine expression in the lungs that were similar to those in the 750-PFU WT group (Figure 4), inoculation with Y17H virus at the normalized 375,000-PFU dose induced upregulation of 80% (289 vs. 161) and 104% (629 vs. 308) more genes at 2 DPI and 5 DPI, respectively (Figure 5 B,C). Although this suggested that Y17H virus might less effectively counteract host-cell antiviral responses, virus dosage effects may also have contributed to the increased gene expression. Therefore, we analyzed the number of shared and unique genes upregulated after infection with WT and Y17H viruses. At 5 DPI, 90% of the genes upregulated in the 750-PFU WT group were also upregulated in the 375,000-PFU Y17H group (Figure 5C). In contrast, only 44% of the genes upregulated in the 375,000-PFU Y17H group were also upregulated in the 750-PFU WT group. A total of 386 genes were uniquely upregulated in the Y17H virus–infected groups at 5 DPI, whereas only 29 were uniquely upregulated in the WT virus–infected group. Thus, Y17H virus appeared to trigger host-cell responses more robustly than did WT virus. Gene enrichment analysis showed that the genes most differentially upregulated were involved in the IFN pathway, the inflammatory response, and/or the antiviral state (Figure 5D). Transcription profiles of genes in the IFN pathway were largely similar for the 750-PFU WT and 375,000-PFU Y17H groups (Figure 5E).

**Figure 5.**
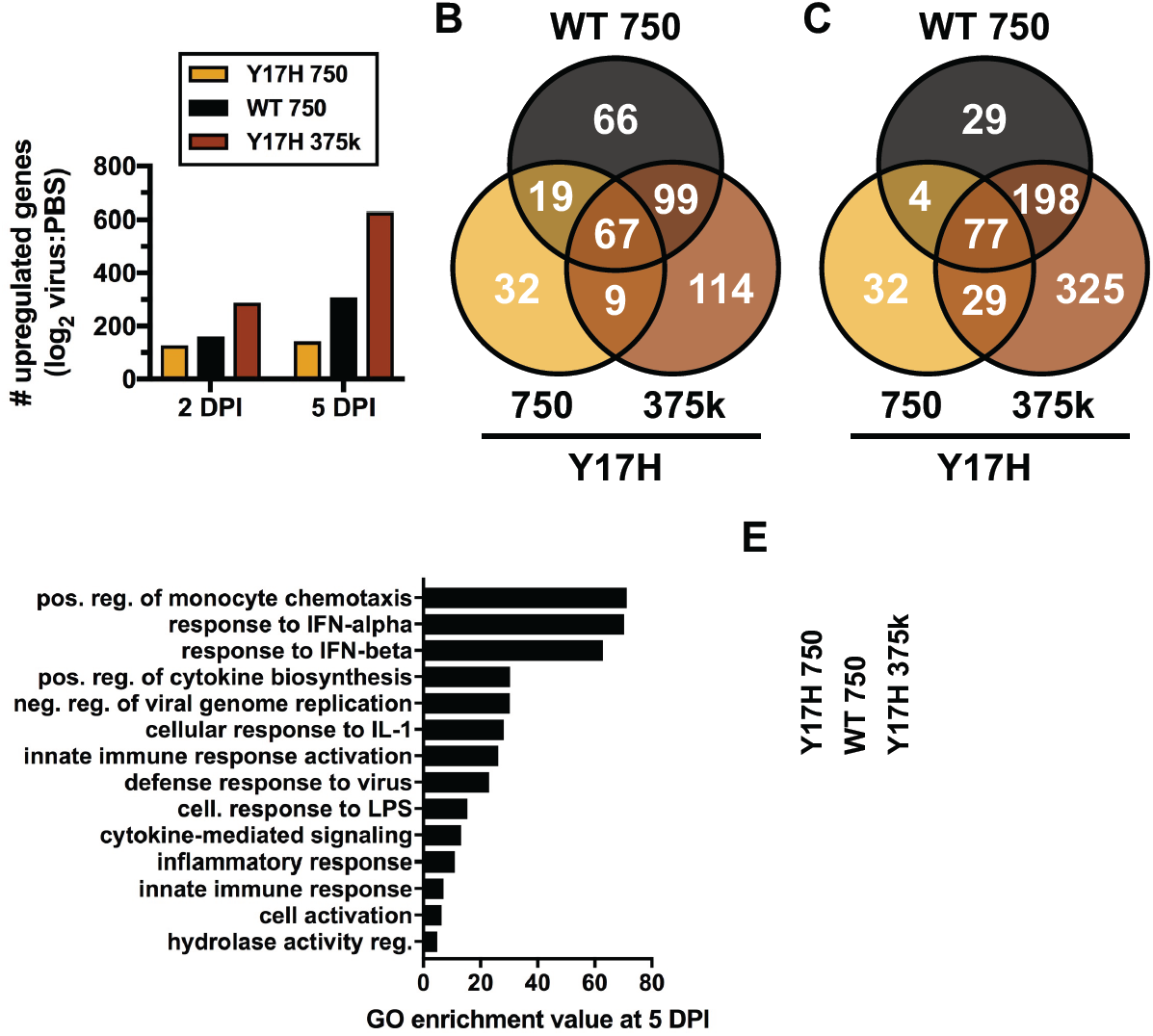
Differential gene expression in DBA/2 mouse lungs after infection. Mice were inoculated intranasally with PBS, WT virus (750 PFU), or Y17H virus (750 or 375,000 PFU). At 2 and 5 DPI, lungs were collected for analysis of RNA expression by Affymetrix (n = 3). **(A)** Numbers of upregulated genes (log2 ratio of virus/PBS > 0.5). **(B,C)** Venn diagrams of upregulated genes. **(D)** Gene Ontology enrichment analysis in the Biological Process category for differentially expressed genes common to all three groups (*P* values < 0.05). **(E)** Heat map of genes involved in the interferon pathways. The log2 ratio of the mean of the virus group to the mean of the PBS group is shown.

To investigate further the effect of the HA-Y17H mutation on type I IFN signaling, we performed Western blot analyses by using antibodies to STAT1, phosphorylated STAT1 (pSTAT1), influenza M1 (as a control for infectivity), and β-actin (Figure 6A). Comparing the 375,000-PFU Y17H and 750-PFU WT groups at 5 DPI, we found that pSTAT1 expression in the Y17H group was 2-fold higher than in the WT group (Figure 6D), but both viruses had similar M1 expression. Overall, Y17H virus induced the upregulation of more host genes and a higher level of pSTAT1 responses in murine lungs than did WT virus when mice were inoculated with Y17H at a dose 500-fold higher than that used for WT virus. The higher dose was necessary to boost the growth of Y17H virus to a level comparable to that of WT virus.

**Figure 6.**
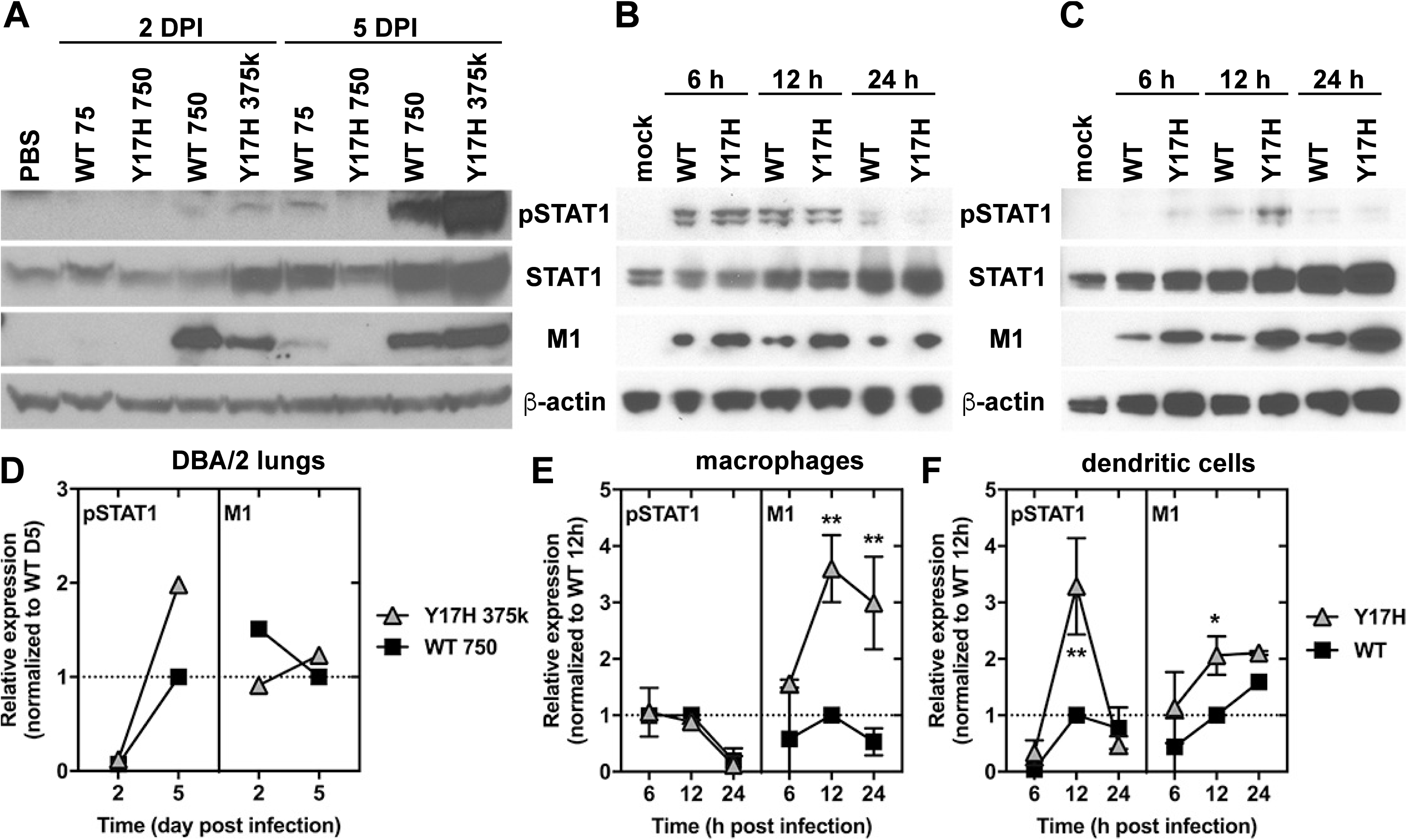
Protein expression by Western blot. **(A)** Mice were inoculated intranasally with PBS, WT virus (75 or 750 PFU), or Y17H virus (750 or 375,000 PFU). At 2 and 5 DPI, lungs were collected for Western blot analysis of protein expression (n = 3). **(B,C)** Bone marrow– derived macrophages (B) or dendritic cells (C) were infected at an MOI of 3 PFU/cell. After 6, 12, or 24 h, protein was harvested for Western blot analysis. SDS-PAGE gels were labeled with antibodies specific for pSTAT1, STAT1, A/H1N1 influenza virus M1, or b-actin. **(D–F)** Quantitation of relative protein expression of pSTAT1 (left panel) and M1 (right panel) by ImageJ. In panel D, the results for the WT 750 PFU and Y17H 375,000 PFU groups from DBA/2 mouse lungs were normalized to WT expression at 5 DPI. In panels E and F, the results for WT and Y17H virus infection (at equivalent MOIs in macrophages and dendritic cells) were normalized to WT expression at 12 h post infection. Values for WT virus (black squares) and Y17H virus (gray triangles) represents the mean (± SD) for a single gel in panel A and for two gels in panels B and C. Significant differences, compared to WT, were determined by multiple Student’s *t-* tests: * *P* < 0.5, \*\**P* < 0.01.

### Type I IFN responses are enhanced in Y17H virus–infected dendritic cells

Dendritic cells and macrophages recognize IAV and other pathogens via a subset of intracellular receptors, such as Toll-like receptors 7/8 (TLR 7/8) and retinoic inducible gene (RIG)-I helicase, and they release high levels of antiviral mediators such as type I IFN and other inflammatory cytokines/chemokines(57). This process leads to more immune cells being recruited to the site of the infection and to the initiation of an adaptive immune response. As the previous analyses of type I IFN responses in murine lungs required a 500-fold higher dose of Y17H virus to normalize the level of infection, we wished to investigate these responses to infection at an equal MOI in the immune cells most likely to display potential differences.

We derived DCs and macrophages from bone marrow of DBA2/J mice by using standard protocols (47). After 6–7 days of differentiation, we infected cells with WT or Y17H virus at an MOI of 3 PFU/cell. In macrophages, M1 expression was more than 3-fold higher in Y17H virus– infected cells at 12–24 h post infection (Figure 6B,E), consistent with macrophages supporting productive infection by Y17H virus but not WT virus in single-step growth-curve experiments (Figure 3I). However, pSTAT1 expression in infected macrophages was similar for Y17H and WT viruses between 6 and 24 h post infection (Figure 6E). In DCs, pSTAT1 responses were highest at 12 h post infection, at which time pSTAT1 expression in Y17H virus–infected cells was 3.3-fold higher than in WT virus–infected cells (Figure 6C,F). Thus, pSTAT1 expression in DCs, but not in macrophages, was elevated by infection with Y17H virus, as compared to infection with WT virus.

We also measured the induction of inflammatory mediators in bone marrow–derived DCs and macrophages infected with WT and Y17H viruses at an MOI of 3 PFU/cell. Infection with either virus resulted in increased expression of IFNβ, CXCL10, IL6, and TNFα at 24 h post infection (Figure 7A–D). Compared to WT virus, Y17H virus induced 2.2-fold higher expression of IFNβ (*P* < 0.0001) and 2.3-fold higher expression of CXCL10 (*P* <0.001) in DCs but not in macrophages. In DCs, NP was expressed at a level 1.13-fold higher during Y17H virus infection (*P* = 0.0313) at 4 h post infection (Figure 7H). At 6 and 12 h post infection, DCs infected with Y17H virus also had higher levels of vRNA, cRNA, and mRNA transcripts than did DCs infected with WT virus (Figure 7E–G). Overall, compared to WT virus, Y17H virus infected DCs, transcribed viral genes, and translated viral proteins at higher levels early in infection, resulting in increased induction of protective responses, including elevated expression of pSTAT1 and the inflammatory mediators IFNβ and CXCL10.

**Figure 7.**
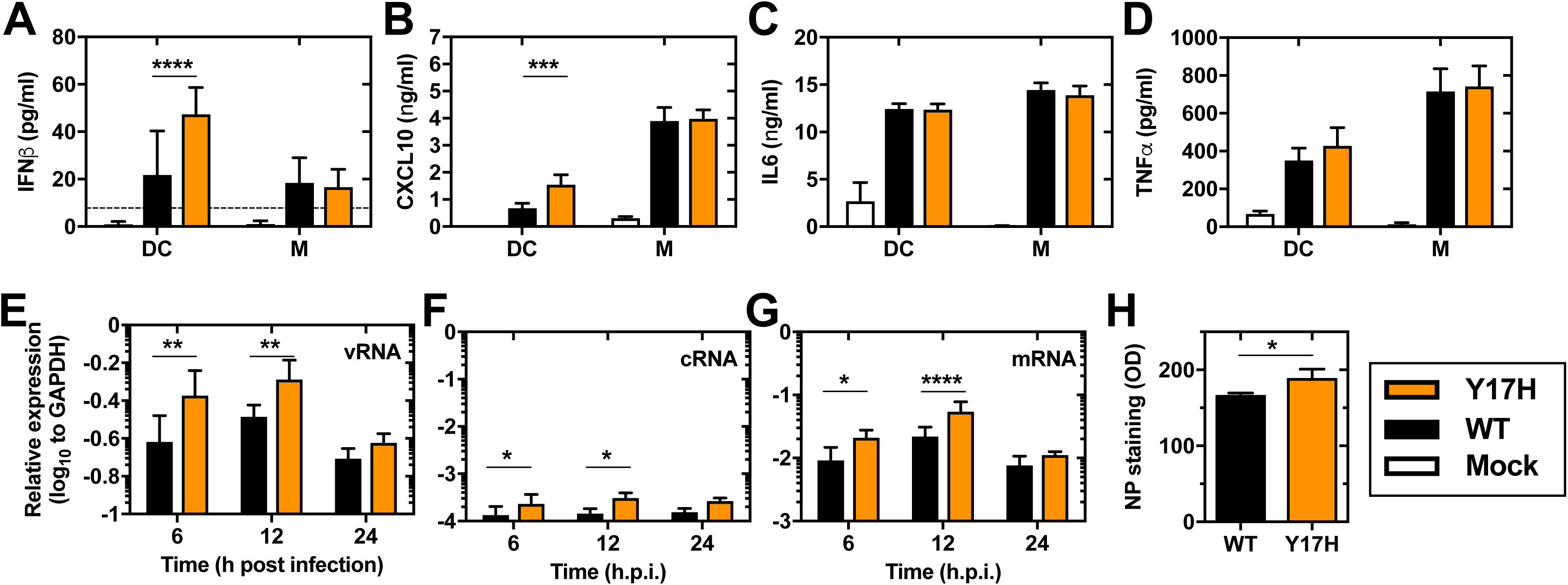
Host responses in bone-marrow derived dendritic cells (DC) and macrophages (M) from DBA2/J mice. Cells were infected with WT and Y17H viruses (at an MOI of 3) or left untreated (Mock). Three experiments were performed independently, with each having two or three replicates. **(A–D)** Cytokine levels were measured in the supernatants by ELISA at 24 h post infection. **(E,F)** Immunoblotting of dendritic cell (E) and macrophage (F) lysates at 6, 12, and 24 h post infection. **(G)** Dendritic cells were stained with an anti-NP antibody 4 h after infection and imaged in a plate reader. **(H)** Viral transcript levels in dendritic cells were measured by real-time RT-PCR. Statistical analysis was performed using one-way ANOVA followed by Tukey’s post hoc tests (A–D) or Student’s *t*-tests (E–H) separately for each day: * *P* < 0.05, ** *P* < 0.01, *** *P* < 0.001, **** *P* < 0.0001.

To investigate if WT and Y17H viruses differentially induced Rig-I like receptors (RLR) and TLR signaling, we infected bone marrow–derived DCs from WT, mice lacking the mitochondrial antiviral-signaling protein (MAVS), and mice lacking the two major adaptor proteins MyD88 and TRIF. After 24 hours of infection, we collected supernatants and measured induction of the cytokines IFNβ, IL6, and TNFα (Figure 8). Consistent with previous results, IFNβ expression was 32% higher in Y17H-infected BMDCs from WT mice than those infected with WT virus (Figure 8A). MAVS is the adaptor protein required for retinoic acid-inducible gene I (RIG-I)-mediated sensing of IAV RNA (58, 59). Bone marrow-derived DCs from mice lacking MAVS were unable to produce IFNβ after infection with either virus, but these cells did induce expression of IL6 and TNFα (Figure 8). The effects were similar for infection with both WT and Y17H virus. These data confirm RIG-I-mediated sensing of IAV was required for interferon signaling in DCs and that both viruses stimulate the RIG-I/MAVS pathway to trigger the type I IFN response in DCs. In bone marrow-derived DCs from mice lacking MyD88 and TRIF, IFNβ expression was reduced compared to IFNβ expression in DCs from WT mice. In these cells, expression of IL6 and TNFα was eliminated, showing the MyD88/TRIF signaling pathway was required to induce IL6 and TNFα inflammatory cytokines. Overall, the results suggest that WT and Y17H are sensed by the same pathway in DCs and that the increased type I IFN response observed during Y17H infection is due to increased replication by the mutant virus.

**Figure 8.**
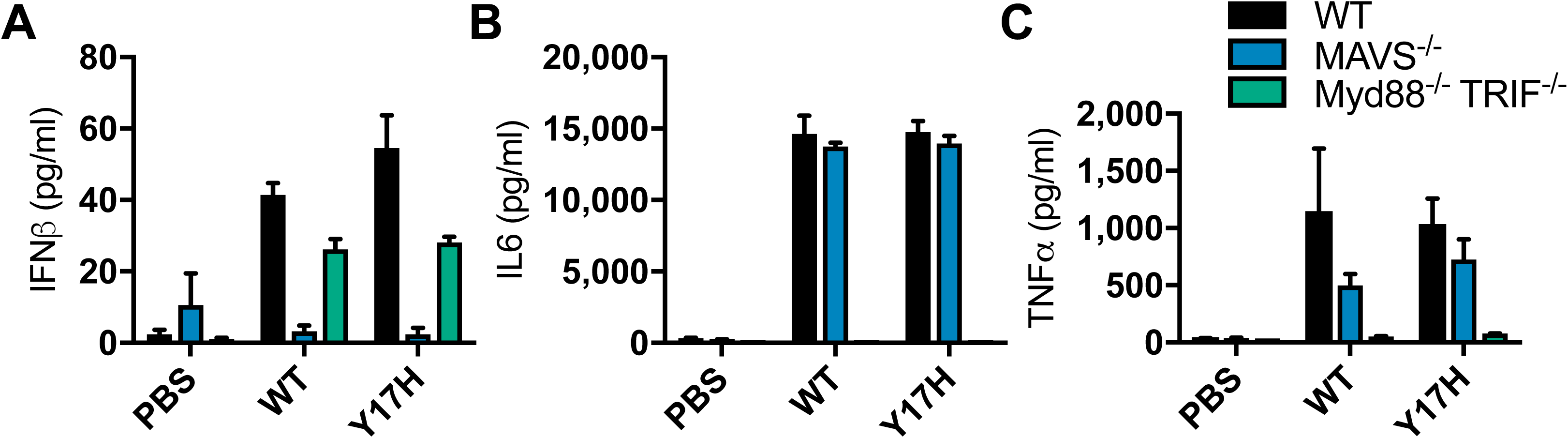
Cytokine production in BMDCs from WT and knock-out C57Bl6 mice. BMDCs from WT (black), MAVS^-/-^(blue), and Myd88^-/-^ TRIF^-/-^ (green) mice were infected with WT and Y17H viruses at an MOI of 3 PFU/cell. Supernatants were collected 24 hours after infection so that the levels of expression of IFNβ (A), IL6 (B), and TNFα (C), could be measured by ELISA. Displayed values are mean (± SD) for 3 mice.

## Discussion

We have investigated several potential mechanisms by which a destabilizing mutation in the HA protein might attenuate the replication of A/H1N1/2009 virus, reduce its infectivity (as demonstrated by an increased ID_50_), and diminish its pathogenesis in mice. The extracellular pH in the lungs of DBA/2 mice ranged from 6.9 to 7.5, a near-neutral range in which changes in HA stability did not affect the persistence of A/H1N1/2009 infectivity. The HA stability altered the virus tropism in cell culture. A destabilizing HA mutation boosted virus replication in single-step growth-curve experiments in MDCK, A549, and Raw264.7 cells but reduced multistep replication in murine airway mNEC and mTEC cultures. In DBA/2 mice, a destabilizing HA mutation substantially reduced virus replication, virus spread in lungs, the severity of infection, and cellular infiltration into BALF. Increasing the inoculation dose of the destabilized HA-Y17H mutant virus 500-fold, as compared to the dose of WT virus, boosted the replication, pulmonary spread, severity of infection, and cellular infiltration to levels comparable to those seen with WT virus. Normalized infection with the destabilized mutant triggered the upregulation of more host genes and increased type I IFN responses and cytokine expression in DBA/2 mouse lungs and bone marrow–derived DCs. Collectively, the data show that HA destabilization attenuates virulence in mice not by decreasing the extracellular virion stability but by enhancing early viral transcription and translation in DCs, which results in stronger activation of antiviral responses, including type I IFN.

Extracellular environments with acidic pH (pH <6.6), warmer temperature, and/or higher salinity decrease the environmental persistence of IAVs (56, 60). Environmental stability can also be decreased by destabilizing HA mutations (41), i.e., those that increase the pH at which the HA protein becomes activated to undergo irreversible structural changes (16, 30). In the present study, the A/H1N1/2009 virus containing a destabilizing HA-Y17H mutation (activation pH: 6.0) had reduced environmental stability over 72 h at pH 6.4, but not at pH 7.0. From this, we hypothesize that such a destabilizing HA mutation will attenuate influenza virus replication in acidic environments but not at near-neutral pH. The respiratory pH in the lungs of DBA/2 mice ranged from 6.9 to 7.5. In previous work, the nasal pH and tracheal pH in mice were reported to range from 6.5 to 7.0 and from 6.7 to 7.5, respectively (61–63). Therefore, we conclude that extracellular inactivation is unlikely to be the cause of A/H1N1/2009-HA-Y17H attenuation in mice. The HA-Y17H mutation may decrease the environmental persistence of A/H1N1/2009 in the human nasal cavity, the pH of which is reported to range from 5.3 to 7.0 in healthy adults and from 5.5 to 6.7 in healthy children (64–67).

The HA-Y17H mutation studied here increases the HA activation pH of A/H1N1/2009 to pH 6.0, most likely enabling quicker virion entry into early endosomes. The wild-type HA protein is activated at pH 5.5, most likely delaying virion entry until the progression into late endosomes. Earlier entry by the Y17H mutant virus is consistent with the accelerated replication kinetics of the virus in single-step growth-curve experiments in MDCK, A549, and Raw264.7 cells and bone marrow–derived DCs. Entry into early endosomes via a relatively unstable HA protein such as HA-Y17H may also enable the virus to avoid functional interactions with IFTM2 and IFITM3 proteins in late endosomes. For reverse-genetics viruses containing the six internal genes of A/H1N1/PR8 and two surface genes, less stable HA proteins from avian influenza viruses were shown to confer increased infectivity in MDCK cells, and the viruses were less sensitive to antiviral IFN responses (28). If these effects contribute to the increased pathogenicity of H7 and H9 viruses in humans, as has been suggested (28), the effect may be lost for seasonal IAVs. On the background of A/H1N1/2009, any early replication advantage by the less stable HA-Y17H protein in single-step growth was lost during multistep replication in MDCK, mNEC, and mTEC cells. In previous publications, destabilizing HA mutations have also resulted in reduced virus growth in DBA/2 mice, ferrets, and human airway epithelium (15, 68). Although the impact of HA destabilization on virulence in humans is unknown, the HA-Y17H mutation was previously associated with a loss of pandemic potential in A/H1N1/2009 (15).

Wild-type A/TN/1-560/2009 (H1N1) virus with an activation pH of 5.5 was replication incompetent in macrophages, whereas the destabilized Y17H mutant with an activation pH of 6.0 produced infectious virions. Similarly, the closely related virus A/CA/04/09 (H1N1) wild-type (activation pH: 5.4) was replication incompetent in Raw264.7 cells, whereas a 7+1 reassortant encoding a less stable H5 HA protein (pH 5.9) produced infectious virions (27). In the present study, gain-of-function replication in macrophages did not correlate with altered antiviral IFN responses or with increased virus replication and pathogenicity in mice. Thus, expanded tropism does not necessarily result in gain-of-function pathogenicity.

Dendritic cells often elicit immunity by capturing and phagocytosing influenza-infected airway cells (69, 70). Additionally, IAVs have been shown to directly infect primary human myeloid DCs, human blood monocyte-derived DCs, mouse splenic DCs, and mouse bone marrow–derived DCs (71–80), although these infections do not necessarily produce infectious virions (80). A/H1N1/09 wild-type virus was previously shown to induce weak cytokine responses in DCs but to be highly sensitive to the antiviral activities of IFNs (73). In the present study, the destabilizing HA-Y17H mutation increased early infection and type I IFN responses in mouse bone marrow–derived DCs, but not in macrophages, and amplified IFN responses in mice were associated with attenuated virus replication and pathogenesis. Thus, the present work shows that the destabilization of the A/H1N1/2009 HA protein to an activation pH of 6.0, a relatively high value typically associated with avian influenza viruses, helps elicit stronger type I IFN responses by accelerating infection in DCs.

We have discovered that HA stability influences IAV replication and pathogenesis in mice by altering the virus infection kinetics and type I IFN responses in DCs. This provides a link between a fundamental property of the viral surface glycoprotein, namely, acid-induced activation of the HA protein, and the ability of the virus to evade antiviral host responses. Further studies are needed to assess the impact of HA stability on the infection kinetics in DCs of other IAV strains and the resulting antiviral responses, especially for highly pathogenic avian influenza viruses that have HA proteins with relatively high activation pH values of similar to the Y17H virus studied here. Future investigations should also investigate the impact of HA stability on IFN responses in pigs, ferrets, and humans. In these studies, it will be important to assess the relative importance of HA stabilization in dampening IFN responses, as has been demonstrated in the present work by infection of DCs vis-a-vis HA stabilization potentially increasing IFN responses by interactions with IFITM2 and IFITM3 proteins in late endosomes (28). Overall, we hypothesize that narrow ranges of HA activation pH support robust replication by a particular IAV in a specific host. The restrictions on the HA activation pH should be considered when developing antiviral drugs and vaccines, especially those that target the more highly conserved stalk region. It should be noted that resistance mutations that alter HA stability without reducing fitness in cell culture, such as the HA-Y17H mutation discussed here, may greatly compromise virus replication in vivo. Thus, antiviral drug and broadly reactive vaccine candidates that target the HA stalk but are susceptible to HA stability mutations in cell culture should not be readily dismissed if such changes compromise the in vivo fitness of the virus.

## Acknowledgments

We thank Teneema Kuriakose for helping with BMDM and BMDC preparation and helpful discussions. We thank Geoffrey Neale for help with the microarray analyses. We thank Richard J. Webby, Thomas P. Fabrizio, and Subrata Barman for providing the IAV reverse genetics system. We thank Keith Laycock of for revising the manuscript. We thank the following facilities at St. Jude Children’s Research Hospital: the Animal Imaging Center, the Animal Resources Center, the Department of Scientific Editing, Hartwell Center Affymetrix, Hartwell Center DNA Sequencing & Genotyping, and the Veterinary Pathology Core Laboratory. This work was funded, in part, by the National Institute of Allergy and Infectious Diseases under Centers of Excellence for Influenza Research and Surveillance (CEIRS) contracts nos. HHSN266200700005C and HHSN272201400006C, by St. Jude Children’s Research Hospital, and by the American Lebanese Syrian Associated Charities (ALSAC).

---

**Figure S1. Results for two pairs of groups inoculated with different virus doses with the aim of normalizing the lung viral loads and weight changes.** In the lower dosing scheme, mice were inoculated with 75 PFU of WT virus or 750 PFU of Y17H virus. In the higher dosing scheme, mice were inoculated with 375,000 PFU of Y17H virus or 750 PFU of WT virus. **(A,B)** Lung tissue titers at 2 and 5 days post infection (DPI) for groups of DBA/2 mice inoculated with the lower (A) and higher (B) dosing schemes. Values are the mean (± SD) for three replicates. **(C,D)** Changes in starting weight for the lower (C) and higher (D) dosing schemes. Values are the combined mean (± SD) for two experiments with a total of 10 mice. Black squares: WT virus; orange triangles: Y17H virus. *P* values were determined for compared pairs by Student’s *t*-test. ns = not significant (*P* > 0.05).

**Figure S2. Microscopic analyses of lung sections.** Mice were inoculated intranasally with PBS, WT virus (750 PFU), or Y17H virus (750 or 375,000 PFU). At 5 DPI, lungs were collected and sections were stained and photographed. **(A)** The extent of active infection (red) and cleared infection (yellow) was visually traced by a blinded histopathologist. **(B)** Hematoxylin and eosin (H&E) staining of lung sections (20× magnification). **(C)** Immunohistochemical staining (IHC) of infected lungs, using a polyclonal antibody against the NP protein.

